# Transcriptional and cell type profiles of cortical brain regions showing ultradian cortisol rhythm dependent responses to emotional face stimulation

**DOI:** 10.1101/2022.01.05.475032

**Authors:** Philippe C Habets, Konstantinos Kalafatakis, Oleh Dzyubachyk, Steven J.A. van der Werff, Arlin Keo, Jamini Thakrar, Ahmed Mahfouz, Alberto M Pereira, Georgina M Russell, Stafford L Lightman, Onno C Meijer

## Abstract

The characteristic endogenous circadian rhythm of plasma glucocorticoid concentrations is made up from an underlying ultradian pulsatile secretory pattern. Recent evidence has indicated that this ultradian cortisol pulsatility is crucial for normal emotional response in man. In this study, we investigate the anatomical transcriptional and cell type signature of brain regions sensitive to a loss of ultradian rhythmicity in the context of emotional processing. We combine human cell type and transcriptomic atlas data of high spatial resolution with functional magnetic resonance imaging (fMRI) data. We show that the loss of cortisol ultradian rhythm alters emotional processing response in cortical brain areas that are characterized by transcriptional and cellular profiles of GABAergic function. We find that two previously identified key components of rapid non-genomic GC signaling – the ANXA1 gene and retrograde endocannabinoid signaling – show top differential expression and the most significant enrichment. Our results further indicate that specific cell types, including a specific NPY-expressing GABAergic neuronal cell type, and specific G protein signaling cascades underly the cerebral effects of a loss of ultradian cortisol rhythm. Our results provide a biological mechanistic underpinning of our fMRI findings, indicating specific cell types and cascades as a target for manipulation in future experimental studies.

## Introduction

Glucocorticoids (GCs) are a class of mammalian hormones known for their pleotropic effects across different bodily systems, such as metabolism, fluid homeostasis, immune and stress system responsivity, as well as brain function. The immunomodulatory capacity of these hormones has been utilised in clinical therapeutics for more than half a century.^1^ The underlying mechanisms through which GCs mediate such a diversity of biological processes remain a topic of intensive investigation. Recent evidence indicates that biorhythmicity might be of great importance. GCs exhibit a circadian rhythm, with high hormonal levels being secreted just prior to and during the active part of the day. The circadian rhythm is superimposed on an underlying ultradian rhythm of more frequent episodes of GC secretion (i.e., hormonal pulses).^2^ The brain is exposed to these hormonal pulses and has developed mechanisms able to perceive them and translate them to cellular, genomic and non-genomic events.^3^ Thus, GC pulsatility might regulate various physiological−, neural−, and glial processes, under baseline and stressful conditions, and hormonal dysrhythmicity could be associated with cognitive and behavioural disorders.^4^

We designed and conducted a randomised, double-blind, placebo-controlled, crossover study to assess functional relevance of GC pulsatility for human brain circuitry. We used a human model of adrenal insufficiency (metyrapone-induced suppression of GC endogenous biosynthesis),^5^ in which GC deficiency was exogenously replaced via two different, pump-mediated subcutaneous infusion methods: one mimicking the normal adrenal function under baseline conditions (resembling the normal circadian and underlying ultradian, pulsatile rhythm) and another lacking GC ultradian pulsatility. The cumulative dosage of the infused hydrocortisone was equal for both methods (20mg/day). Exposure of the human brain to the same emotional stimuli (fearful, happy, and sad faces) provokes a differential response from corticolimbic regions of the right hemisphere, involved in emotional processing, depending on the mode (i.e., presence or absence of ultradian rhythmicity) of GC replacement.^6^ These functional magnetic resonance imaging (MRI) findings provide evidence that ultradian GC rhythm could be critical in regulating neural dynamics in human, but, at the same time, they raise the question why these particular brain regions show to be sensitive to changes in GC rhythmicity, while other brain regions did not.

In the current work, we approached this question from a transcriptional and cell type point of view: we investigated the relationship between differential GC rhythm-dependent brain activation in the fMRI data and anatomically patterned transcriptional and cell type profiles. To do this we utilized available data from the Allen Human Brain Atlas (AHBA).^7^ This is an anatomically comprehensive transcriptional brain atlas sampled from a number of carefully selected, clinically unremarkable donor brains, produced by a combination of histology-guided fine neuroanatomical molecular profiling and microarray-assisted mapping of gene expression data into MRI coordinate space. The AHBA provides an unparalleled high-resolution genome-wide map of transcript distribution and the ability to analyze genes underlying the function of specific brain regions.^8–12^

In this context, we combined the functional MRI results of our study with available AHBA data to investigate which genes of the AHBA donor brains were differentially expressed in the GC rhythm-sensitive cortical brain areas (as specified by our functional MRI study) in comparison with the remaining cortical areas, thus specifying an anatomical transcriptomic signature of GC rhythm-sensitive cortical brain areas. We utilized gene ontology, pathway and protein-protein interaction databases to look for enrichment of functions (i.e., relate gene expression profiles of brain GC rhythm sensitiveness to enrichment of specific brain cell functionality). We also utilized AHBA neuronal cell type databases to scale up the signature of cortical brain GC rhythm sensitiveness from a transcript to a cell type level. In the latter case, if specific human-verified neuronal cell types are discovered to be enriched in the areas that show GC rhythm-responsivity, this could trigger a selective, preclinical, experimental investigation on the relationship between these human neuronal cell types and GC rhythmicity.

## Results

### Differential expression analysis

The brain regions that showed significant variations in the BOLD signal responses to emotional stimulation (exposure to emotionally valenced faces) between the pulsatile and the non-pulsatile group were matched with brain coordinates of the AHBA samples from the left cortex, as depicted in Figure 1 (see Methods section for details). Note that although lateralization in function is well described, it has been established in multiple studies that there are no statistically significant transcriptional hemispheric differences in adult brain.^7,13^ This indicates that post-transcriptional factors constitute hemispheric differences in function, while left and right hemispheres are similar on a gene transcription level. Because only two donor brains have been sampled bilaterally, while all six donor brains include samples from the left hemisphere, we maximized spatial resolution by mapping the fMRI effects of the right hemisphere to cortical AHBA sample coordinates of the left hemisphere by inverting the x-axis coordinates in MNI-152 space (Figure 1A). After selection of cortical samples, based on their inclusion in the cortical Desikan parcellation atlas,^14^ 61 left cortical samples were mapped to the differentially responsive brain areas (regions corresponding to either “mask A” or “mask B” in Figure 1A), and 1224 left cortical samples were selected as control samples (Figure 1B and Table 1).

**Figure 1.**
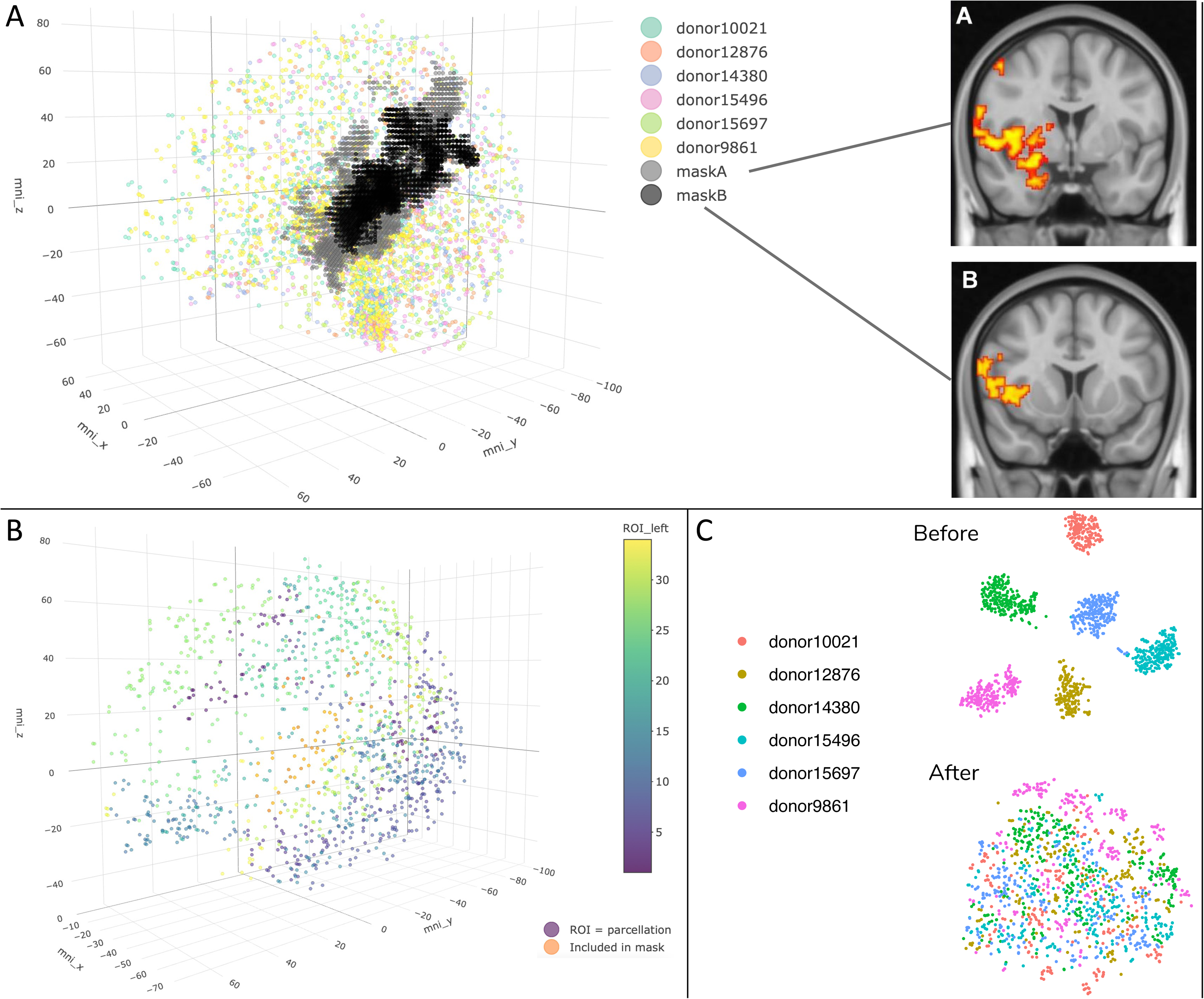
**A.** 3702 AHBA brain samples plotted with left-mapped brain regions that show differential responsiveness to GC pulsatility in the same three-dimensional (MNI-152) space. “Mask A” and “mask B” (black and grey dots in the 3D plot) correspond to the left-mapped versions of the brain regions that show differential responsiveness to GC pulsatility in the right hemisphere (2D images A & B). **B.** 1285 left cortical samples included in the differential expression analysis. Samples are colored depending on the Desikan parcellation they are mapped to (34 parcellations in total). Samples included in the mirrored fMRI mask coordinates (A and B from Figure 1) are presented in orange. **C.** tSNE on the 1285 included left cortical samples before (left) and after (right) applying our normalization strategy (see Methods section for details).

**Table 1.** Overview of samples from the six AHBA donors in the differentially GC-rhythm responsive area and the control regions.

|  | Donor<br>10021 | Donor<br>12876 | Donor<br>14380 | Donor<br>15496 | Donor<br>15697 | Donor<br>9861 | All<br>donors |
| --- | --- | --- | --- | --- | --- | --- | --- |
| Differentially<br>responsive<br>area | 6 | 10 | 10 | 13 | 8 | 14 | 61 |
| Control<br>region | 169 | 168 | 249 | 209 | 222 | 207 | 1224 |

Despite the gene expression normalization procedures performed by the AHBA,^15^ it has been established that large inter-individual differences in gene expression remain in the AHBA samples.^16^ We too find that for the 1285 left cortical samples used in our differential expression analysis, samples from the same brain have more similar gene expression levels. To account for these between-donor effects, and additionally any between-sample effects in our differential expression analysis, we applied both within-sample and across-sample normalization strategies (see Methods section).^16^ We visualize the efficacy of these strategies by running a t-distributed Stochastic Neighbor Embedding (tSNE) on all 1285 included cortical samples before and after preprocessing steps (Figure 1C).

After all preprocessing steps and probe selection criteria were applied, differential expression analysis was performed for 10015 gene transcripts, comparing 61 AHBA samples (cases) with 1224 AHBA samples (controls). This resulted in 304 genes that showed significant differential expression after correcting for false discovery rate (FDR), with the significancy threshold set at Benjamini-Hochberg corrected p < 0.05. 223 genes showed a differentially higher expression, and 81 genes showed a differentially lower expression. The top 25 of differentially expressed genes are plotted in Figure 2A (for a full list see Supplementary Table 1).

**Figure 2.**
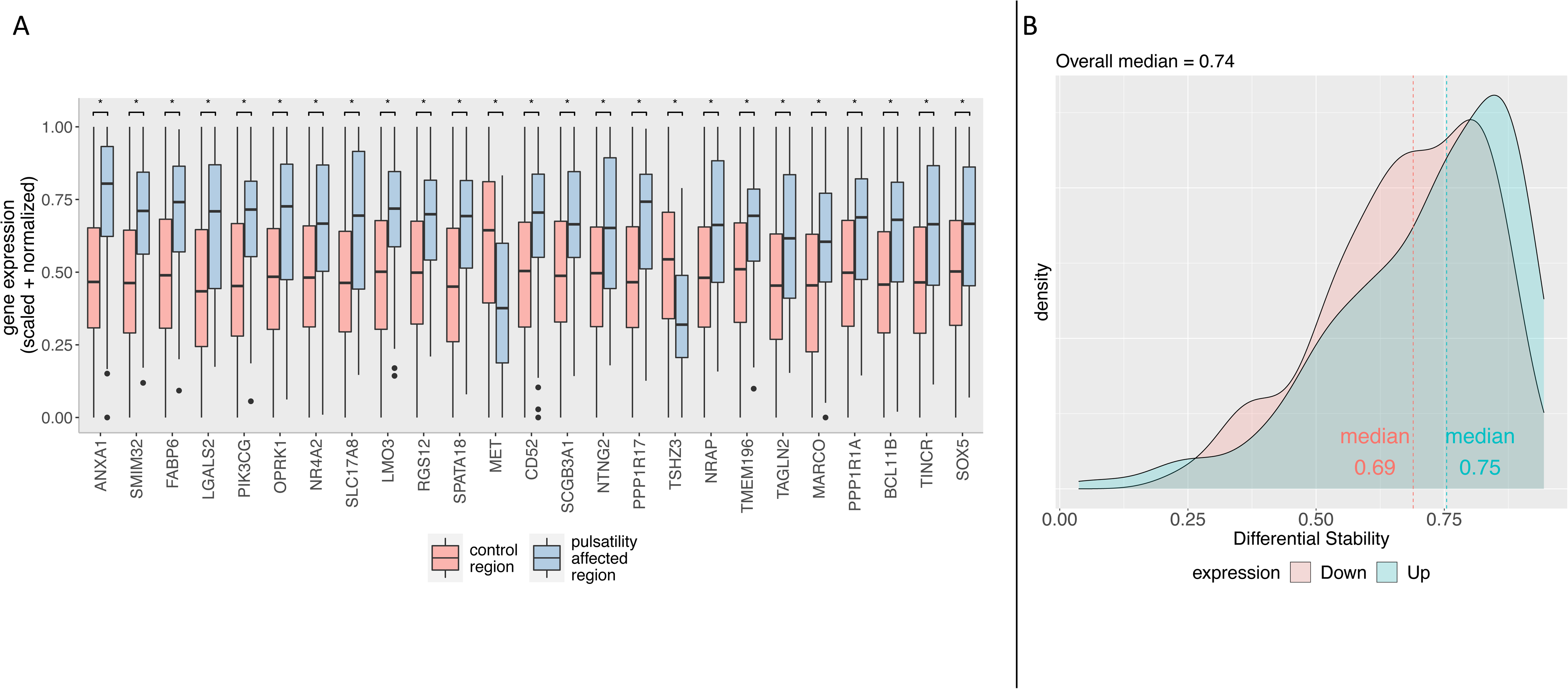
**A**. Boxplot of top 25 differentially expressed genes, ordered by significance from left to right (increasing p-value from left to right). The bottom and top hinges correspond to the first and third quartiles, with the median shown in the interquartile range (IQR). Whiskers extend to the smallest and largest values within a range of 1.5*IQR from the bottom or top hinge. Values outside the 1.5*IQR range are plotted as individual outliers. All plotted results have FDR corrected p < 0.05 (*). **B**. Density plot of differential stability values, plotted separately for higher and lower differentially expressed genes.

Although possible donor-driven effects were mitigated by extensive normalization strategies, we additionally tested the possibility of incomparable transcriptomic signatures between donor brains (for example due to age, gender, ethnicity etc.) that might invalidate the differential expression analysis. We did this by using the differential stability (DS) metric: a correlation-based measure for the consistency of a gene’s differential expression pattern across brain structures.^17^ We used the fact that gene expression patterning across brain structures was assessed for reproducibility in all six AHBA donor brains in previous works.^18,19^ By cross-referencing our differentially expressed genes list with these previous results (Supplementary Table 2 of the study by Hawrylycz et al.^18^), we find that our differentially expressed genes show high median differential stability in comparison to other genes (median = 0.74 versus median = 0.53, p < 0.001). This indicates that the identified genes have reproducible gene expression patterns across all donor brains, regardless of sex, age and other donor related factors (Figure 2B).

### Differential gene expression in pulsatility-sensitive brain regions shows neuronal specificity and enrichment for retrograde endocannabinoid signaling

We analyzed the differentially expressed genes for enrichment of gene ontology (GO) terms (including GO terms relating to cellular components, biological processes, and molecular functions) and Kyoto Encyclopedia of Genes and Genomes (KEGG) pathway categories (Figure 3). Using all 223 genes that showed higher differential expression in the pulsatility-responsive brain regions, we found significant enrichment for 37 different GO terms related to cellular components and biological processes (FDR corrected p < 0.05, see Supplementary Table 2). Most notably, terms relating to neurons (e.g. neuron part, neuron development, neuron projection) and intercellular communication (e.g. signaling, cell communication, synapse, response to stimulus) were found to be enriched, confirming neuronal specificity of differential gene expression in pulsatility-responsive brain regions. Significant enrichment for KEGG pathways (FDR corrected p < 0.05) was found for the categories ‘retrograde endocannabinoid signaling’, ‘glutamatergic synapse’, ‘GABAergic synapse’ and ‘morphine addiction’ (see Figure 3 and Table S3). In the 81 genes with lower differential expression, none of the positive hits in the GO term enrichment analysis showed statistically significance after FDR correction (Supplementary Table 4). Enrichment analysis for KEGG pathways in these 81 Genes yielded one positive hit but reached no statistically significance (Supplementary Table 5).

**Figure 3.**
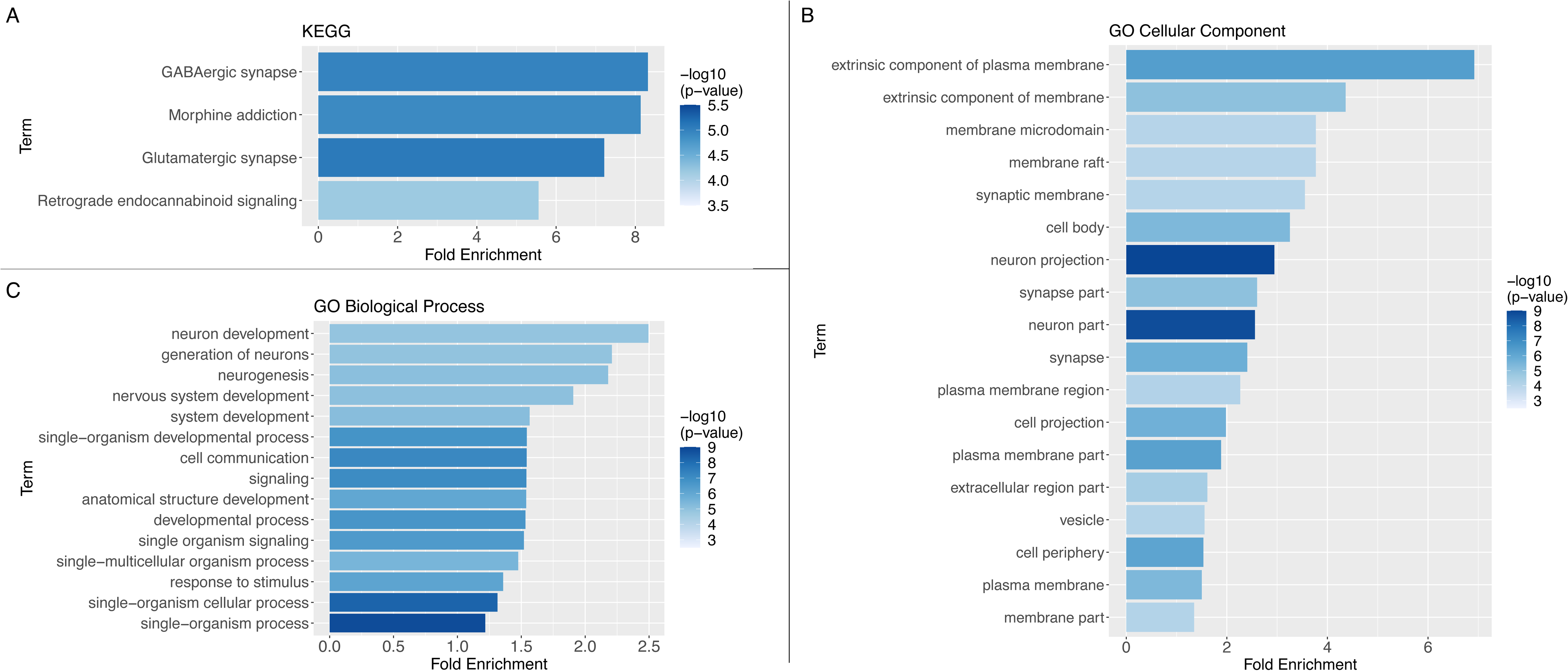
Functional enrichment analysis results for **A)** KEGG pathways, **B)** GO categories relating to cellular components and **C)** GO categories relating to biological processes. Only term enrichments with Bonferroni-adjusted p < 0.05 are shown. Terms are ordered according to fold enrichment relative to chance, with colours indicating the −log_10_() transformed nominal p-value (higher −log_10_() value means lower p-value)

### Differentially expressed genes in pulsatility-sensitive cortical brain regions show enrichment for transcriptomic signatures of stress-related psychiatric disease

Using two recent brain transcriptomic studies defining brain transcriptomic signatures for several psychiatric disorders based on several tissue types,^20,21^ we tested for enrichment of our higher and lower differentially expressed genes in these transcriptomic signatures (see Table S7 for all results). In one study including both MDD and PTSD brain samples,^20^ we found enrichment of both MDD and PTSD genes in our 223 higher differentially expressed genes (Table S7). Using a second study including multiple psychiatric diseases,^21^ our differentially higher expressed genes were enriched for genes showing higher differential expression in autism spectrum disorder (ASD), bipolar disorder (BD) and alcoholic abuse disorder (AAD). In our 81 differentially lower expressed genes, we found enrichment for genes showing differentially lower expression in ASD, schizophrenia (SCZ) and BD (Table S7). In further validation of our differential expression results in relation to stress-related disorders, we performed additional enrichment analysis using the PheWeb database: a dataset based on genome-wide associations for EHR-derived diagnoses in the UK Biobank.^22^ We found the most enriched category (having the highest odds ratio) to be ‘Acute reaction to stress’ (see Figure S2).

### Protein-protein interaction enrichment analysis reveals association of pulsatility-responsive brain regions with particular G_i_α signaling events

Using the list of 223 differentially expressed genes that show higher expression in the pulsatility-responsive brain regions, protein-protein interaction enrichment of the proteins encoded by those genes was analyzed using multiple databases (see Methods section) to plot a network containing all proteins (encoded by our differentially expressed gene list) that have documented interactions with at least one other protein in the list. Next, using the MCODE algorithm^23^ for detection of densely connected network components within this network, we found two densely connected networks. One densely connected network consisted of ADRA1B, GNAO1, GNG2, GNG4, GNB2, GNB4 and PIK3CG. Subsequent pathway and process enrichment analysis of this MCODE component indicates functional enrichment for G beta-gamma signaling through PIK3Kgamma, more generic G-protein beta-gamma signaling and cholinergic synapse (Figure 4). The other densely connected network consisted of protein-protein interactions between NPY, ANXA1, SST, PTGER3, OPRK1 and CXCL2, and showed functional enrichment for Gα_i_ signaling events, and (more generic) G protein-coupled receptor (GPCR) ligand binding, as well as a subclass of GPCRs, the rhodopsin-like receptors. These results further support the notion that GC rhythm alterations act directly on the brain and will have most effect in brain regions that, on an anatomical transcriptional patterning level, show to be enriched for genes related to particular GPCR functionality. The same analysis performed on the set of genes with lower differential expression did not result in any densely connected networks showing significant enrichment.

**Figure 4.**
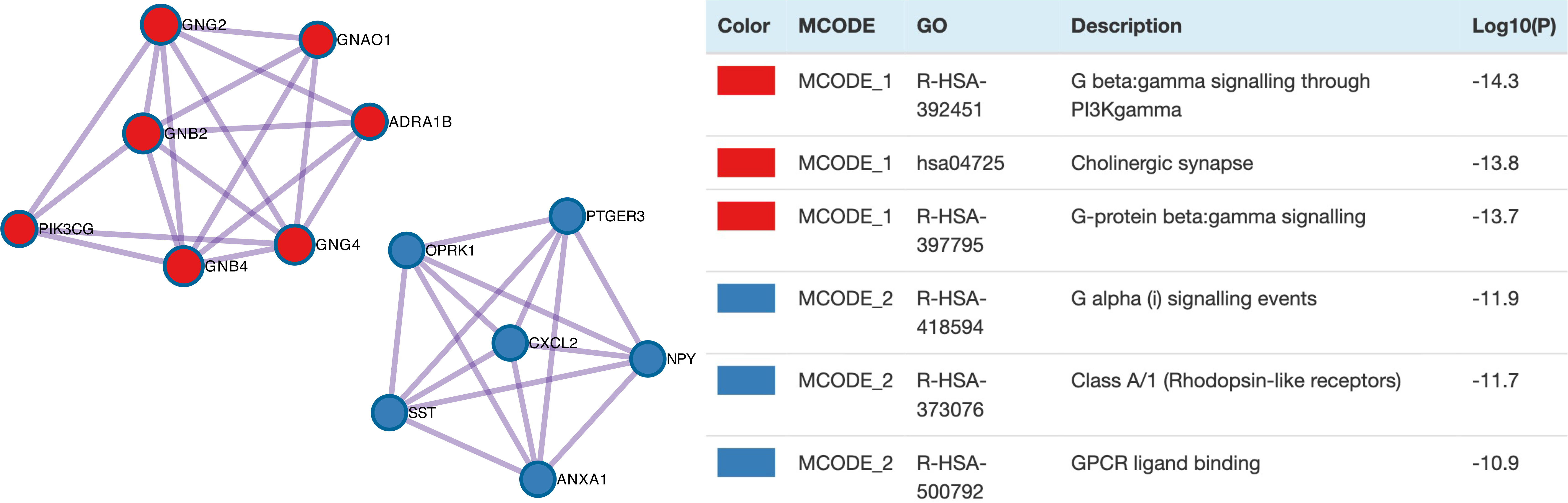
Densely connected networks of protein-protein interactions and their enrichment categories. GPCR: G protein-coupled receptor.

### Cortical cell type analysis reveals exclusive enrichment of GABAergic neurons and differentially expressed genes specific to certain cell types

Using the 223 higher differentially expressed genes, human cortical cell type enrichment analysis showed an exclusive enrichment for GABAergic cell types. Specifically, the three cell types found to be enriched (Figure 5) were the GABAergic SP8-expressing interneurons (belonging to a cluster of neurons expressing PAX6 and TNFAIP8L3); EGFEM1P-expressing interneurons (belonging to the cluster of neurons expressing VIP and PENK); and the QRFPR-expressing interneurons (belonging to the cluster of neurons expressing SST and GXYLT2). Using the 81 lower differentially expressed genes showed no enrichment for any of the neuronal cell types (Figure 5).

**Figure 5.**
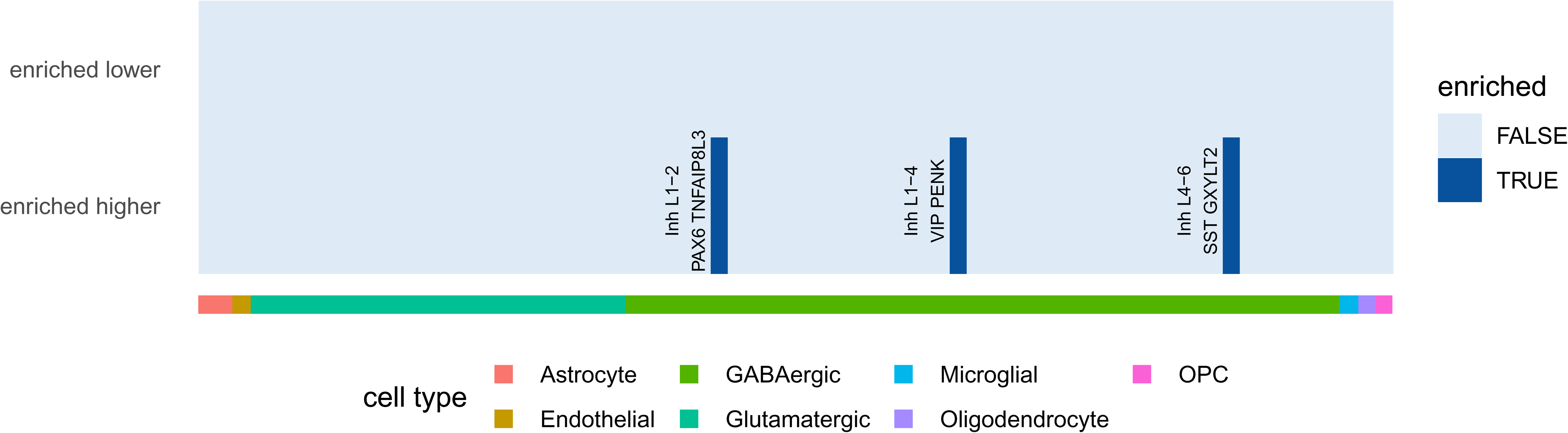
Results of human cortical cell type enrichment analysis reveal four inhibitory neuronal cell types for the differentially higher expressed genes (DEG higher). For the lower expressed genes (DEG lower) no neuronal cell types were found to be enriched.

For the 223 higher differentially expressed genes, cross-referencing of single marker genes, specific and sufficient for the classification of a single cortical neuronal cell type, yielded a match with the GABAergic NPY-expressing interneurons. When taking all marker genes for this cell type into account, the NPY-expressing neurons failed to reach significance in our hypergeometric enrichment analysis, as not all marker genes were present in our differentially higher expressed gene set. However, in the cell type dataset, NPY gene expression was found to be highly specific for a single inhibitory cell type (Figure S1).^24^ Even though the NPY-expressing neuronal cell type failed to reach significance in the hypergeometric test (p = 0.265), we thus consider the significantly higher expression of NPY in the examined brain regions an indication of an enrichment of the specific NPY-expressing GABAergic neurons in those regions.

To further look on a cell type level at the MCODE component genes in the protein-protein interaction analysis, which also contains NPY, we used the gene expression levels by cell type from a recently added single nucleus RNA sequencing cluster-based cell type dataset, consisting of 120 distinguished human cortical cell types (see Methods section).^25^ The results are plotted in Figure 6, and again show a specific GABAergic neuronal cell type (“Inh L6 SST NPY”) that shows highly specific expression of the NPY gene. The same plot including the top-50 higher differentially expressed genes and both cortisol receptors (GR and MR) is shown as Figure S1.

**Figure 6.**
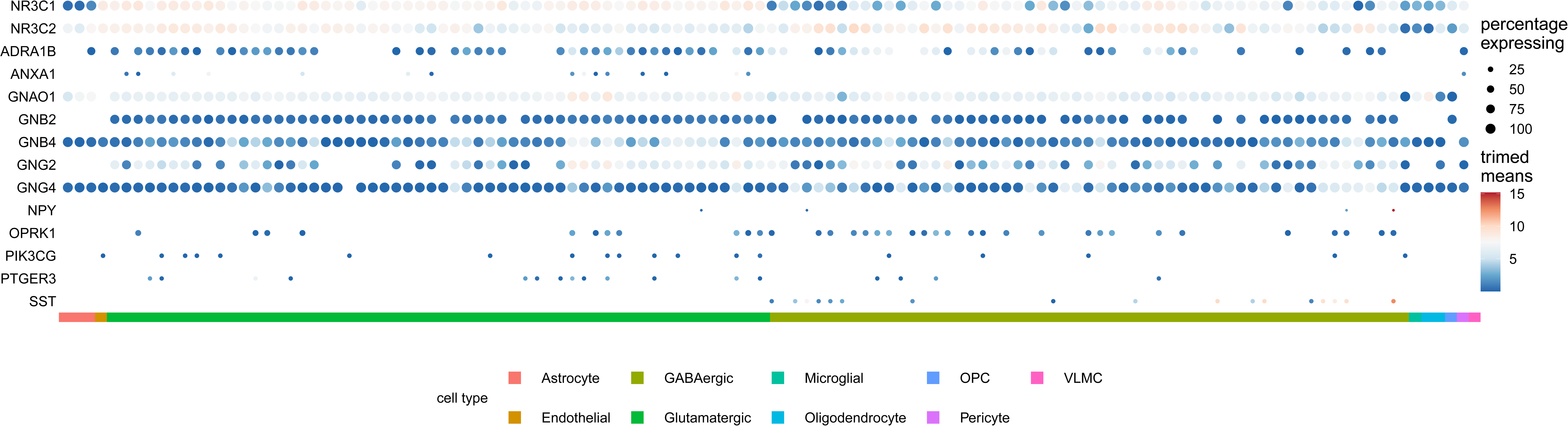
Dot plot of trimmed mean gene expression per cell type for GR, MR, and the identified MCODE cluster, and their specificity (percentage of cell types expressing them). Genes with a trimmed mean expression of 0 in all cell types (meaning they show expression in less than 25% of cells for each cell type) are omitted from the plot (see Methods section for details). The red arrow points to high mean expression (dark red colored dot) of NPY very specific for a particular SST-expressing GABAergic neuronal cell type.

In addition to the specific human cortical neuronal cell type datasets, that include a vast majority of cortical neuronal cell types, we also looked at enrichment in a larger cross-laboratory, rodent-derived dataset containing putative marker genes for different cell types in the brain, not specific to either cortical regions or neuronal subpopulations.^26^ We found a significant enrichment for marker genes of microglia, Purkinje cells and serotonergic cells (Supplementary Table 6).

## Discussion

In this study we investigate the transcriptional and cell type patterning of brain regions that are sensitive to cortisol pulsatilty. Strikingly, both the top significant differentially expressed gene (ANXA1) and the most significantly enriched KEGG pathway term (retrograde endocannabinoid signaling) have been identified before as key components of rapid non-genomic GC signaling.^27–30^

The endocannabinoid system has emerged as an important regulator of some of the rapid, non-genomic glucocorticoid effects in the brain.^28–30^ The mechanism involves the glucocorticoid-mediated activation of membrane associated variants of the GRs at the target brain cells (mainly postsynaptic sites) to induce endocannabinoid synthesis and/or local release, causing retrograde cannabinoid type I receptor-mediated modulation of the presynaptic neuronal activity. This mechanism has been described as responsible for (i) the hormonal negative feedback regulation of excitatory synaptic inputs to hypothalamic (paraventricular nucleus) neuroendocrine cells,^28^ and (ii) the long-lasting suppression of spontaneous inhibitory synaptic inputs^29^ as well as facilitation of excitatory inputs to the basolateral amygdala principal neurons induced by glucocorticoids in the acute stress setting,^31^ while (iii) a similar involvement of this glucocorticoid-induced mechanism has been also described for the attenuation of inhibitory transmission in prelimbic cortex, again under stressful conditions.^32^ Our data suggest that cortical brain regions with the capacity of recruiting the retrograde endocannabinoid signaling pathway may be more sensitive to the characteristics of the ultradian glucocorticoid rhythm, i.e., able to convert changes in glucocorticoid pulsatility into different neurobiological effects (in our case differential neural activation in response to the same emotional stimuli).

The ANXA1 finding is also of great interest as it has been implicated as a facilitator in the rapid non-genomic inhibitory feedback effects of endogenous glucocorticoids on ACTH release.^27^ Although these findings were limited to the folliculo-stellate cells in the anterior pituitary gland, the ANXA1 gene has also been implicated in the neuroprotective and anti-inflammatory role of microglia.^33^ Although sparse ANXA1 gene expression has been demonstrated in microglia of human brains,^33^ the extensive cortical cell type data based on single nucleus RNA sequencing of multiple cortical regions of the human brain^25,34^ that we used here, shows ANXA1 gene expression to be mostly apparent in excitatory neuronal cells (Figure S1). In this these data, ANXA1 does not show any pronounced expression (trimmed means > 0) in the microglial cell types of the human cortical cell type data (Figure S1). ANXA1 is however listed as a microglial marker gene in the NeuroExpresso rodent-based data.

ANXA1 is also part of the defined densely connected network of protein-protein interactions that drives enrichment of terms related to GPCR function, more specifically Gα_i_ signaling events. This enrichment can be interpreted as an indication of i) the involvement of a specific (set of) cortisol activated GPCR(s) associated with G alpha i proteins, ii) membrane GR/MR having a hitherto unknown association with Gα_i_ signaling events, iii) cross-talk between GR/MR signaling and signaling cascades involving these G protein species. For example, this could mean that a cortisol pulse is linked to an inhibitory transmission effect (Gα_i_).^35^ Additionally, the interaction network showed enrichment of G-protein beta and gamma subunits and linked this to acetylcholine receptor signaling. It was previously demonstrated that intracellular calcium signaling via the mAChR3 subtype depends on the beta 2, gamma2 and gamma 4 subunits that are present in the MCODE-1 network.^36^ This suggests a link between calcium regulation and the sensitivity to cortisol pulses, perhaps involving annexin A1.^37^

The results of our cell type analysis show exclusive enrichment for GABAergic neurons, with NPY gene expression enrichment indicating possible enrichment of a specific SST-NPY expressing GABAergic interneuron. This is of interest because evidence has consistently implicated (SST expressing) GABAergic interneuron dysfunction in MDD (Major Depressive Disorder) pathology.^38,39^ Both hypercortisolemia and circadian rhythm alterations have been related to MDD subtypes and MDD pathology in general.^40^ Our results additionally seem to point to the importance of ultradian rhythm disturbances in the process of GABAergic interneuron dysfunction in MDD pathology. In our differentially expressed genes, we did in fact find enrichment of an MDD brain transcriptomic signature as defined by one recent study.^20^ However, we did not find an MDD brain transcriptomic signature enrichment when using the transcriptomic signature defined by another study.^21^

Although anatomical cell type enrichment in our defined brain regions appears to be exclusive for GABAergic neurons in the human cortical cell type data, inspection of the cell type specific expression shows excitatory neuron specificity for some differentially expressed genes as well. Specifically, ANXA1 and NR4A2 show expression predominantly in glutamatergic neuronal cell types (see Figure S1). This implies that, although there might be an important role for GABAergic signaling in pulsatility sensitivity, the Gα_i_ and Gα_q_ signaling events implicated in the differential brain activational response to cortisol rhythm changes are not specific to GABAergic cell types but occur in glutamatergic neurons as well.

No microglial enrichment was found in the human cortical cell type analysis, but it was found in NeuroExpresso-based results. The enrichment analysis using the NeuroExpresso data, however, is more difficult to interpret, as the marker gene data used here are rodent derived, not cortex specific and is not comparable to the standard of tissue processing and measuring protocols from the human cell type data (i.e. mainly microarray based, different labs with different procedures were used).^25^ Also, while only six marker genes were included for microglial cells in the human cortical cell type data,^24^ the NeuroExpresso datasets lists over a hundred unique marker genes for microglia (see Supplementary Table 6). Depending on the specificity of those genes, an abundance of marker genes could lead to an increase in false positive results. Yet, there is considerable evidence for a role of microglia in the stress response and transcriptomic dysregulation of microglia, although mostly in the context of overactivation in chronic stress.^41,42^

### Limitations and strengths

It is important to note that we only describe anatomical transcriptomic patterns, based on healthy donor brains, and do not investigate putative brain transcriptional effects in the brains of the participants of the pulsatility study. The aim of our current study was to investigate the anatomical transcriptional and cell type patterning of pulsatility sensitive brain regions. Notably, this is unrelated to any local transcriptional effects that loss of ultradian cortisol rhythm might have on specific brain regions, and does not include any post-transcriptional factors that can explain observed lateralization in function.

The AHBA provides the most detailed dataset for examining spatial distribution of human brain transcriptomics to date but is limited to six donor brains. Another differential expression analysis with the same spatial resolution would not be possible in other publicly available data at this time. Importantly, it should be noted that to mitigate any donor-driven bias, and to control for any possible incomparability of gene expression levels due to donor-specific differences (e.g. age, gender, ethnicity) we 1) did thoroughly correct for any donor-driven gene expression bias, 2) used a regression based method that allows for repeated sampling from the same subjects, and 3) checked the comparability of gene expression patterning of differentially expressed genes across donor brains using the differential stability criterion. The outcome of these control measurements (see Figure 1C and 2B) indicate an unbiased differential expression analysis and reproducible results across all 6 donor brains, regardless of demographic or biological differences of donors. Although we estimate the sample size (i.e. n = 1285) coming from these six donors to be sufficient for the scope of our current analyses, including more donor brains with the same spatial resolution likely would have further improved the generalizability of the current results to brains with a more substantial diversity of donor characteristics.

We define a cortical brain region as ‘pulsatility sensitive’, i.e. showing a differential pattern of neural activation between physiological pulsatile and non-pulsatile groups, on the basis of a task-based fMRI study. It is therefore possible that additional cortical regions could be defined as ‘pulsatility sensitive’ in a different task-based setting. Our results thus formally define transcriptional and cell type patterns of regions that are pulsatility sensitive in the context of emotional processing. Accordingly, we selected ‘rest-of-cortex’ as control samples in our differential expression analysis because the fMRI results are based on a whole-brain analysis - in the context of emotional processing. Importantly, these fMRI results reflect significant differences in neural activation during emotional processing between both groups (see supplementary figure S4 of Kalafatakis et al).^6^ This means that the statistical maps we used in our analysis are unrelated and independent to mean task activation (i.e., a brain region might show a distinct mean neural response to the task, but might not at all show differential fluctuation of activation during task conditions between groups, and vice versa). Consequently, it would be invalid to select ‘control samples’ only from regions showing significant base neural response to the task. To illustrate, we thresholded a separately calculated statistical map for mean task activation to exactly include all significantly differential responsive regions (meaning that the significantly differential responsive region with the lowest mean task activation is used as a threshold for “task activation”). Using this threshold, virtually the whole brain can be defined as ‘activated by task’. For these reasons, we chose to use “rest-of-cortex” as control samples.

The task-based setting used in our fMRI study, with a validated paradigm for probing emotional processing, has several advantages. First, we found differences in emotional processing on a functional level between pulsatility groups, meaning that the imaging effects correlate to functional effects as well.^6^ Second, by using the context of emotional processing, effects of loss of ultradian rhythm are likely related to brain functionality affected by cortisol signaling related pathologies such as major depressive disorder, posttraumatic stress disorder and other stress related psychiatric disorders.

In fact, we did find enrichment of differentially expressed genes related to transcriptomic brain signatures described in both PTSD and MDD, but also ASD, BD, SCZ and AAD. A difficulty with interpreting these enrichment results is that defined differentially expressed gene sets in both transcriptomic studies used for gene set definition^20,21^ are based on several brain tissues from non-overlapping anatomical origins. None of the tissues included in these studies overlaps with the fMRI mask we used for our differential expression analysis. In this regard it is also important to note that even the same neuronal cell type can have a different transcriptomic profile depending on the cortical region it is embedded in, indicating the possible loss of power using several different non-overlapping tissues.^43^ Another issue is that MDD enrichment was found when using one study by Girgenti et al,^20^ but not when using the MDD gene set as defined by the study by Gandal et al.^21^ This further indicates the difficulty of comparing different brain tissues, especially given a lack of robust marker genes throughout tissue types (which was the case for the data by Girgenti et al). Although the study by Gandal et al lists correlated log2 fold change in differentially expressed genes across disorders, and results from this study that are shared amongst different brain areas and diseases might be more robust, there is only one gene that satisfies the condition of having an FDR-corrected p-value < 0.05 in all five disorders (gene: CRH). This might explain the lack of enrichment of our differential expression results for the ‘all-diseases’ group of genes as defined by the Gandal study.

Cell type enrichment analysis was based on human cortical cell type data, which we consider a strength of the study. However, as these data distinguishes more inhibitory than excitatory neuronal cell types, and lists approximately twice as many combined inhibitory marker genes than excitatory marker genes (386 versus 173), there is a possible bias towards inhibitory cell enrichment. The significantly higher expression of the NPY gene – which is highly specific for a specific SST expressing cluster of inhibitory neuronal cell types –, and the fact that the differentially expressed genes show enrichment for four inhibitory cell types, but no excitatory cell types, do seem to point to the importance of a GABAergic neuronal response to cortisol pulsatility.

As discussed in relation to microglia, the additional cell type enrichment analysis using NeuroExpresso cell type data might not reflect optimal marker gene specificity. For some enriched cell types, NeuroExpresso marker genes also seem to be non-specific upon further inspection (Supplementary Table 5). For example, the enrichment found for ‘serotonergic cells’ is based on the inclusion of two genes (TRH and PTGER3), that are in fact not specific for serotonergic cells. Obviously, serotonin producing cell bodies should be absent or very scarce in cortical cells. Assuming that these markers are not abundant in axonal projections of 5-HT neurons, and given that these ‘marker’ genes are not exclusive for 5-HT neurons, we consider this enrichment call to be a false positive outcome.

The aim of this study was to investigate the anatomical transcriptional and cell type patterning of pulsatility sensitive brain regions. We show that the loss of cortisol ultradian rhythmicity alters emotional processing response in cortical brain areas that are characterized by transcriptional and cellular profiles of GABAergic functioning. Our results indicate that specific cell types and G protein signaling cascades underly the cerebral effects of loss of physiological cortisol rhythm, thus making these cell types and cascades a target for manipulation in future experimental studies.

Overall, in this study, we have identified target genes, signaling pathways and neuronal subtypes that might constitute key players in the physiological response to glucocorticoid pulsatility and its translation to differential biological effects in the human brain.

## Materials and Methods

### Functional MRI study

This was a randomized, double-blind, placebo-controlled crossover study of different modes of hydrocortisone replacement in healthy subjects, registered with the United Kingdom Clinical Research Network (IRAS reference 106181, UKCRN-ID-15236; October 23, 2013). The study followed the CONSORT guidelines for randomized controlled trials. Fifteen right-handed, healthy male volunteers aged 20–33 years were included in the study. The Ethics Committee of the University of Bristol approved the study, and all participants provided informed written consent. More details on the development and validation of the human model of adrenal insufficiency, and the different modes of GC replacement therapy, inclusion and exclusion criteria of the study, recruitment process, quality control and bioethical concerns, randomization, and blinding processes, as well a detailed presentation of all outcome measures recruited (aside functional MRI), can be found elsewhere.^4–6^

### Functional MRI data

The functional image pre-processing steps consisted of (i) brain intensity normalization, (ii) 3D motion correction, (iii) B_0_ unwarping with assistance from the B_0_ fieldmap images, (iv) brain extraction, (v) spatial smoothing, (vi) temporal high pass filtering, and (vii) co-registration of the functional image with a corresponding high-resolution, anatomical, T_1_-weighted image and with MNI152 standard space. Bias field correction has been applied, before removing the non-brain tissue from the high-resolution image. For each individual/ session functional MRI dataset, a regression analysis was performed using a general linear model fitting the temporal evolution corresponding to the paradigm (emotional face presentation). A fraction of the temporal derivative of the blurred original waveform was added to the model. Temporal filtering was also applied. The form of the hemodynamic response function convolution method applied to the basic waveform was the Gamma variate. Three different effects were modelled (original exploratory variables); visual exposure to (i) fearful human faces, (ii) happy human faces and (iii) sad human faces. For the statistical analysis of the functional MRI data acquired during the presentation of emotional faces, we produced individual session/subject level maps of activity, indicating which brain regions were responding to the emotional face recognition (contrasting the baseline, resting state condition). For the comparisons between the GC pulsatile and non-pulsatile groups, whole-brain, group-level analyses were carried out using a mixed effects model. Each group-level analysis produced thresholded z-score brain region clusters highlighting statistically significant variations in the activation pattern between the GC pulsatile and non-pulsatile groups in response to emotional face stimulation. In all cases, corrections for multiple comparisons were performed at the cluster level using Gaussian random field theory (minimum z > 2.3, cluster p threshold < 0.05).^6^

### Transcriptomic atlas

The Allen Human Brain Atlas (AHBA) is a publicly available transcriptional atlas based on microarray measures, using a set of 58692 probes in 3702 samples across brainstem, cerebellum, subcortical and cortical brain structures across six postmortem human brains (five males and one female, age range 24 – 57, African American, Caucasian and Hispanic ethnicities). For limited samples of two donor brains, expression values were also measured by RNA sequencing. All expression data and metadata were downloaded from the AHBA (http://human.brain-map.org) on October 14th, 2019.

### Data analysis

Our data analysis method can be summarized into seven distinct steps (see below). Many of our choices for data handling have been based on the work of Arnatkeviciute et al.^16^ For step two (probe selection) and part of step three (sample selection), MATLAB scripts from their processing pipeline (publicly available at https://github.com/BMHLab/AHBAprocessing/) were adapted and customized for our own analysis (with their approval), using MATLAB version R2020a. For the rest of the analysis steps, except for probe reannotation, the programming language R (version 3.6) has been utilized. All used R packages were installed under R build 3.6. All R code is made publicly available on GitHub at https://github.com/pchabets/fMRI-Transcriptomics-Ultradian-Cortisol.

#### Step 1: Reannotation of probes

Since the probe annotation originally provided by the Allen Institute dates from a decade ago, probes were first reannotated to the latest human genome version and reference sequence using the Re-Annotator pipeline.^44^ Re-Annotator is freely available for download at https://sourceforge.net/projects/reannotator/. The most recent genome and reference sequence were downloaded from the UCSC website on May 20th, 2020. The reannotation step resulted in the selection of 46039 probes annotated to a total of 20200 unique genes for inclusion into further analysis.

#### Step 2: Probe selection

If multiple probes were annotated to the same gene, we selected the best representative probe for that gene. Previous work has shown that selecting probes on the basis of the highest expression (intensity based filtering) improves the mean correlation between microarray and RNA sequencing (RNAseq) measures of gene expressions obtained in the same brain samples, thus improving microarray data reliability.^16^ Therefore, we first selected probes that showed a signal above noise signal in at least 50% of cortical and subcortical samples across all subjects. This resulted in 31977 remaining probes, annotated to 15719 unique genes. Next, for each gene, if multiple probes were annotated to that gene, one single probe was selected by choosing the probe with the highest correlation to the RNAseq measures for the same gene in the same samples (RNA sequencing data available for first two donors. Data were downloaded at http://human.brain-map.org/static/download on October 14th, 2019). To further improve reliability of the differential expression analysis based on microarray probe measurements, we removed probes from the analysis that: a) were annotated to a gene that was not detected by the RNAseq measurement in the same sample, b) showed a low correlation to RNAseq data (Spearman Rho < 0.2). This resulted in 10014 probes selected for 10014 unique genes. Previous work showed that functional enrichment analysis of genes that are removed based on these criteria show no enrichment for genes related to neuronal function.^16^

#### Step3: Sample selection

Two AHBA donor brains were sampled bilaterally, while the other four donor brains were only profiled on the left hemisphere. This was done because no significant interhemispheric transcriptional difference was found in the first two brains. This is in accordance with previous evidence indicating that indeed no statistically significant transcriptomic differences between the left and right hemispheres exist.^7,13^ To maximize spatial coverage, we only included AHBA samples from the left cerebral cortex for analysis. Because the differentially responding brain regions in the considered fMRI data were located on the right hemisphere, we symmetrically flipped the MNI-coordinates of the affected brain regions from right to left, to optimize spatial transcriptomic coverage. Flipping the fMRI mask from right to left is a valid approach in this case, since we are looking at cortical anatomical transcriptional patterning only, and no statistically significant hemispheric difference exists on the mRNA level.

Left hemisphere samples were selected if they could be annotated to the Desikan cortical parcellation atlas, using the AHBA processing pipeline available at https://github.com/BMHLab/AHBAprocessing/.14 In total, 1285 left cortical samples were included for differential gene expression analysis. Next, AHBA samples and fMRI masks were plotted in MNI152 space. Using trilinear interpolation, it was calculated for each AHBA sample whether it could be assigned to an “affected” brain area (meaning falling inside either of the two fMRI thresholded masks) or not. This resulted in 61 samples in the “affected” brain regions versus 1224 samples in the “unaffected” brain regions.

#### Step 4: Normalization of expression values

To verify the possibility that donor-driven effects could bias our differential expression analysis, we plotted the 1285 samples based on their respective gene expression values in a 2D representation using tSNE. This clearly showed clustering of samples by donor brain. We therefore corrected for possible donor-driven effects by using the *RemoveBatchEffect* function from the limma package for R,^45^ treating each donor as a separate batch. Because the limma function uses linear modelling, this correction method can be sensitive to outliers. Therefore, an additional outlier-robust normalization strategy was performed using scaled robust sigmoid (SRS) normalization:

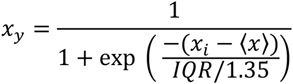

where 〈*x*〉 and IQR represent the median and the inter-quartile range respectively, followed by rescaling to a unit interval of 0-1:^16,46^

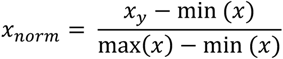

Figure 1C shows the effect of this normalization strategy on the donor-driven clustering of samples. To also account for gene outliers within each sample (within-sample, across-genes normalization), we additionally performed the same SRS and unit-interval scaling procedure within each sample, across all the measured gene expressions.

#### Step 5: Testing significance of differentially expressed genes

Differential gene expression analysis between the 61 “affected” versus 1224 “unaffected” samples was performed using the limma package for R.^39^ Genes were ranked in order of evidence for differential expression by first fitting a linear model to the microarray data with the *lmFit* function, and then using an empirical Bayes method to shrink the probe-wise sample variances towards a common value and to augmenting the degrees of freedom for the individual variances with the *eBayes* function.^45,47^ To further correct for the fact that samples in the “affected” and “unaffected” regions came from the same six donor brains, we included a correlation term for samples coming from the same donor brain using the *duplicateCorrelation* function. We passed this as argument in the *lmFit* function before passing the resulting fit to the *eBayes* function to allow for repeated measures from the same subjects.

False discovery rate correction was performed by using the Benjamini-Hochberg (BH) procedure. BH-corrected p-values of p < 0.05 were considered significant. To check reproducible gene expression patterns of these differentially expressed genes across all donor brains, we cross-referenced publicly available results of calculated differential stability (DS) metrics for the same AHBA data (Supplementary Table 2 of the study by Hawrylycz et al.^18^), plotting the distribution of DS values for the differentially expressed genes (see Figure 2B). Differential stability is a correlation-based measure for the consistency of a gene’s differential expression pattern across brain structures.^17^ A difference in median differential stability of the differentially expressed genes in comparison with other genes was tested using a Mann-Whitney test.

#### Step 6: Functional and protein-protein interaction enrichment analysis

Functional enrichment analysis of differentially expressed genes was performed using the RDAVIDWebService package for R.^48^ Gene ontology terms for biological processes, molecular function and cellular components, as well as pathways described in the KEGG database, were considered. As a background, the transcriptome wide coverage of the AHBA microarray probes was used. Enrichment analysis in the PheWeb database^22^ was performed using the webinterface of Enrichr,^49–51^ with as input the 223 differentially higher expressed genes. We used a hypergeometric test to test for enrichment of gene sets defined as differentially expressed in psychiatric diseases according to two recent transcriptomic signature studies.^20,21^ Data was extracted from DataTable S1 of the Gandal study,^21^ and from the Supplementary Tables 1 and 22 of the Girgenty study.^20^ Genes were included in a disease group if they had an FDR-corrected p-value <= 0.05 for the defined disease, and were divided into ‘higher’ or ‘lower’ differentially expressed according to having a log fold change higher than 0.1 or lower than −0.1 respectively. More stringent criteria resulted in empty gene groups. Inclusion of genes and categories of diseases according to both studies are summarized in Table S7. For protein-protein interaction enrichment analysis and densely connected network discovery, Metascape^52^ was used with the list of 223 differentially higher expressed genes that showed FDR-corrected p < 0.05. Metascape uses the following databases: STRING, BioGrid, OmniPath and InWeb_IM. Only physical interactions in STRING (physical score > 0.132) and BioGrid are used.

#### Step 7: Cell enrichment analysis

To translate our results from the individual gene level to a cell type level that is verified in human, we used the results from a recent Allen Brain Atlas dataset that used samples from human cortical areas (from the middle temporal gyrus) to perform single nucleus RNA sequencing followed by cortical neuronal cell type classification.^24^ This dataset consists of a classification of 75 different GABAergic and glutamatergic cortical neuronal cell types, as well as cortical astrocytes and microglia. We used a hypergeometric test to test for enrichment of certain cortical cell types by concatenating the marker genes for each specific cell type that is listed in Supplementary Table 2 of the paper by Hodge et al.^24^ We used a threshold of at least five different cell markers per cell type, which excluded five cell types (three inhibitory, and two excitatory neuronal cell types: Inh L2-5 VIP SERPINF1, Inh L4-6 SST B3GAT2, Inh L4-5 SST STK32A, Exc L5-6 THEMIS C1QL3, Exc L6 FEZF2 OR2T8). We also cross-referenced our differentially expressed gene set for cell types that showed a single gene to be specific as a cell type marker, as differential expression of such specific single-marker-genes might indicate differential expression of the related cell types – even if the hypergeometric test fails to reach significance because of missing differential expression of other less specific marker genes.

To plot mean gene expression levels in human cortical cell types, we used recently added single nucleus RNA sequencing data, sampled from several locations of human cortical donor brains (middle temporal gyrus, anterior cingulate cortex, primary visual cortex, primary motor cortex, primary somatosensory cortex, primary auditory cortex), available at the AHBA cell type atlas website, listing trimmed mean expression for each gene per distinguished cell type.^25^ This dataset distinguishes a total of 120 different human cortical cell types. Trimmed means are calculated by first log2 transforming gene expression and then calculating the average expression of the middle 50% of the data (data with lowest and highest 25% of expression values removed) independently for each gene and cell type.

Additionally, we used the NeuroExpresso database to perform a similar cell type enrichment analysis for the higher differentially expressed genes.^26^ Genes were converted from rat to human orthologues using the “homologene” package in R, which is a wrapper for the Homologene database by the National Center for Biotechnology Information (NCBI).^53^ Although rodent-derived and not specific for cortical neuronal cell types, this dataset contains more abundant putative marker genes for 20 other cell types, like microglia, oligodendrocytes and astrocytes. The number of included marker genes for each cell type are listed in Supplementary Table 5.

## Acknowledgments

Philippe Habets would like to express his gratitude to Aurina Arnatkevičiūtė for her support and guidance in the field of fMRI transcriptomics, the availability of the MATLAB pipeline for AHBA data, and to the highly insightful and useful work on combining AHBA and imaging data she and others of the Fornito lab have published. Konstantinos Kalafatakis would like to express his gratitude to Bodossaki Foundation (https://www.bodossaki.gr/en/) for supporting the research on modelling biomedical data, especially related to glucocorticoid biorhythmicity. The fMRI data was obtained from studies funded by the Medical Research Council and the Wellcome Trust to Stafford Lightman.

## Supplementary Material

**Figure S1:** Dot plot of trimmed mean gene expression (see Methods section for details) per cell type and their specificity (percentage of cell types expressing them). Genes included are GR, MR, and top-50 genes (higher differentially expressed). Genes with a trimmed mean expression of 0 in all cell types are omitted from the plot.

**Figure S2:** Enrichment analysis results of the PheWeb database using the Enrichr web interface with as input the 223 differentially higher expressed genes from our fMRI-transcriptomics analysis.

**Table S1:** Table of differentially expressed genes (Benjamini-Hochberg corrected p < 0.05) with additional probe and gene data.

**Table S2-3:** Functional enrichment charts using all higher differentially expressed genes, tested for GO terms (molecular function, biological processes, cellular component) and KEGG pathway categories.

**Table S4-5:** Functional enrichment charts using all lower differentially expressed genes, tested for GO terms (molecular function, biological processes, cellular component) and KEGG pathway categories.

**Table S6:** Results of NeuroExpresso cell type enrichment analysis, with marker genes listed.

**Table S7:** Results of enrichment analysis of sets of genes defined as differentially expressed in several psychiatric diseases according to two recent studies.

