## Supplemental Figure 2 for "Transcriptional and cell type profiles of cortical brain regions showing ultradian cortisol rhythm dependent responses to emotional face stimulation"

### PheWeb 2019

Bar Graph | Table | Clustergram | Appyter

Click the bars to sort. Now sorted by combined score ranking.

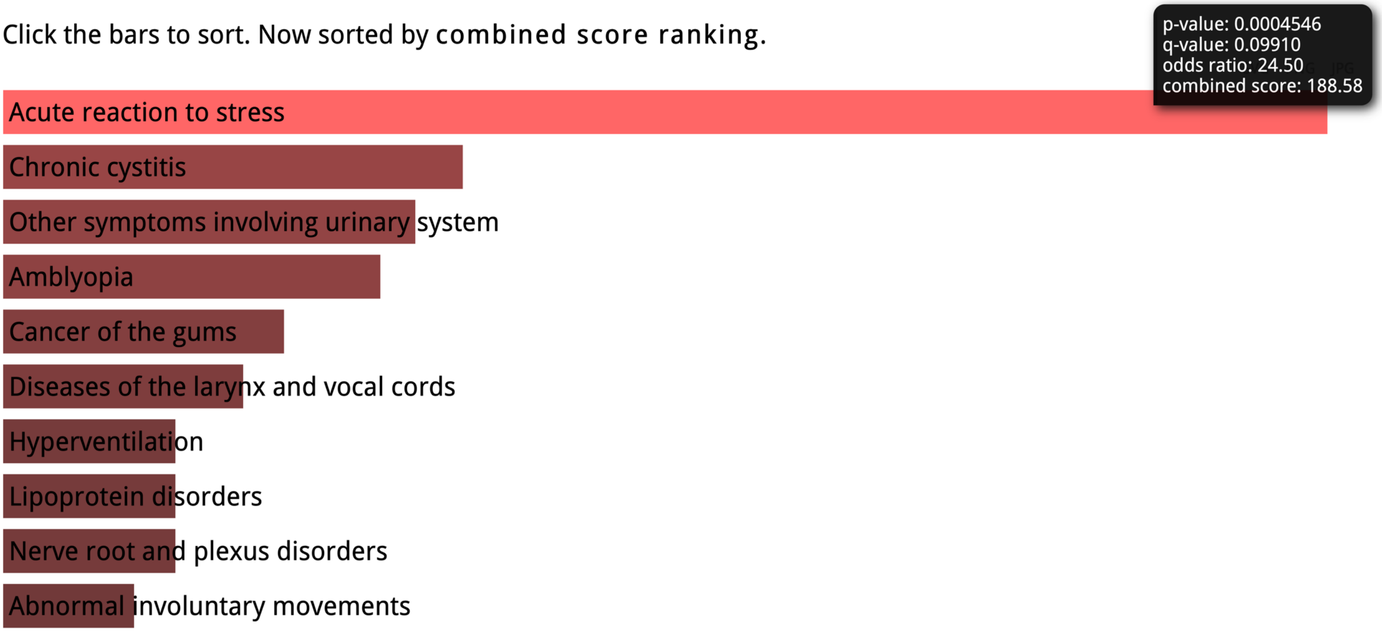
