## Supplementary figures and images for "Transcriptional and cell type profiles of cortical brain regions showing ultradian cortisol rhythm dependent responses to emotional face stimulation"

### Supplemental Figure 1

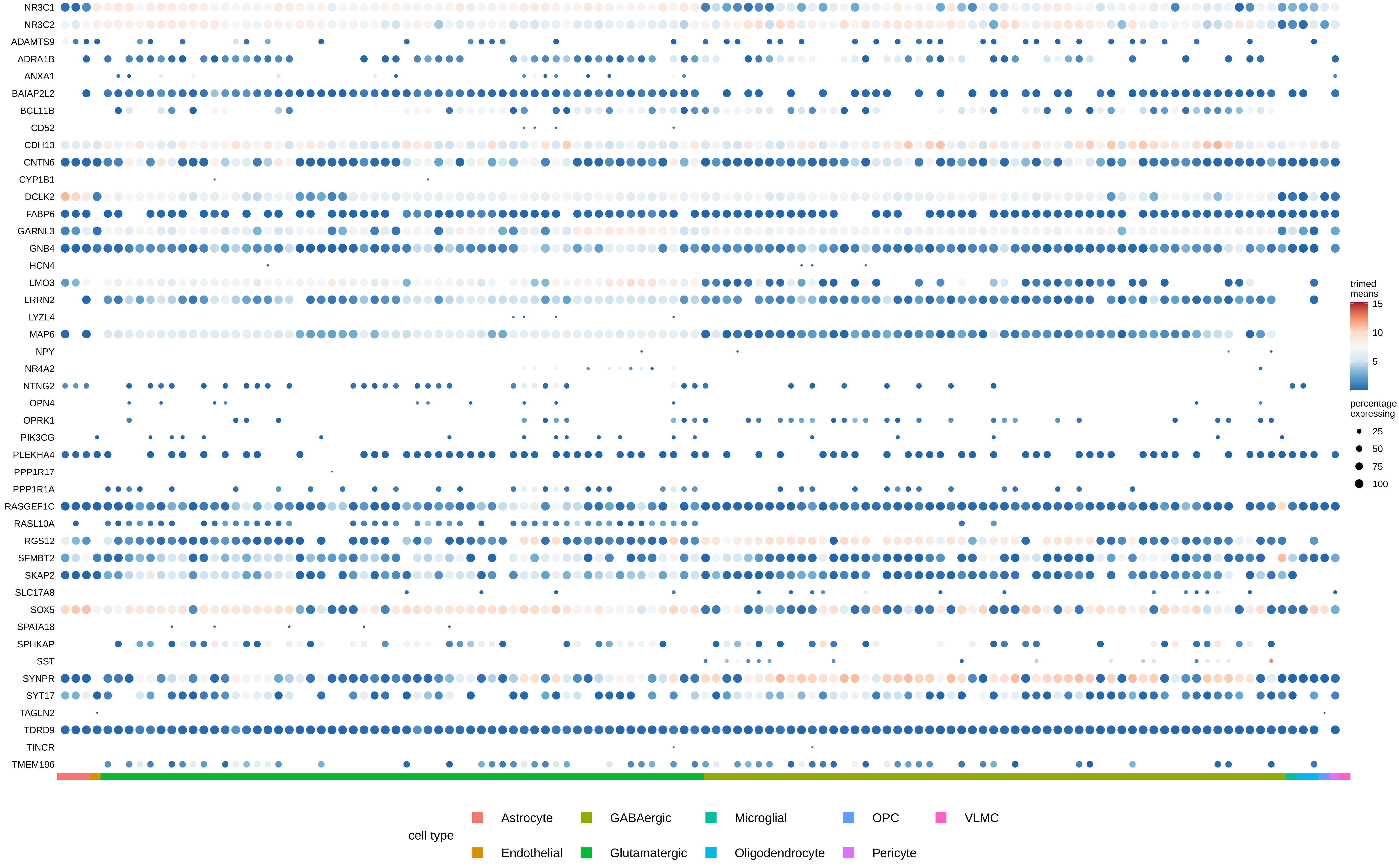
